# Isolation of a novel strain of Lactic Acid Bacteria from traditionally fermented common lime (*Citrus aurantifolia*) of Assam, India and analysis of exopolymeric substances produced by the strain

**DOI:** 10.1101/643650

**Authors:** Nabajyoti Borah, Arindam Barman, Debabrat Baishya

## Abstract

A gram positive, rod shaped and catalase negative strain of Lactic Acid Bacteria was isolated from traditionally fermented common lime (*Citrus aurantifolia*) of Assam, North-East India. Bacterial identification was done by using conventional morphological and biochemical methods as well as advanced molecular technique. Traditionally fermented lime juice was serially diluted on selective culture medium and growth of translucent, ropy bacterial colony was observed in the culture plate. Isolated bacteria were identified up to species level by using ribosomal RNA (rRNA) gene sequencing technique. Based on nucleotide homology and phylogenetic analysis the isolate was found to be a strain of *Lactobacillus delbrueckii*. This is the first report of finding this sub species of Lactic Acid Bacteria in citrus fruit product. The sequence determined in this study has been deposited in the GenBank database with sequential accession number KT198973. The bacterial isolate also produced exopolysaccharide when grown in chemically defined medium. Fourier Transform Infrared Spectroscopy (FTIR) was done for chemical and compositional characterization of partially purified exopolysaccharide.

## Introduction

Lactic acid bacteria (LAB) are one of the most important groups among bacteria due to the “generally recognized as safe (GRAS)” status of this group and their products. The members of this group are mainly characterised for the formation of lactic acid as a final product of the carbohydrate metabolism [1, 2]. The possibility of Lactic acid bacteria in the field of health and nutrition is humongous and this can boast up different sections of traditional treatments. The evidence that LAB can stimulate the immune system is astounding and captivating and opens numerous research areas regarding mechanisms and effective utilization[3]. One of the important aspect of using LAB derived products comprises its cost affectivity as these products can be produced without much difficulty and sophisticated instruments. They have moderately low level of toxicity contrasted with other microbial products [4]. The growing concerns about health and well-being, along with an interest in consuming natural foods, have given probiotics a unique position in the food market. There are lot of documented works with regard to the presence of Lactic acid bacteria in dairy products and for long they are being used extensively as starter culture[5]. However, research had rarely been centered on the microflora found in fermented fruits for the presence of this specific group of bacteria although the microenvironment and nutritive profile of fruits which are rich in nutrients are quite conducive for such bacteria to multiply[6]. Ong et al. reported enterococcus type of LAB in fermented red dragon fruit juice [7]. These bacteria were also been isolated from ripe mulberries and other traditional fruit juices [8, 9].

Nutritious natural products like fruits and vegetables are the normal part of the human diet and are consumed in large quantities by most of the world population. Traditionally, products of plant origin have been viewed as microbiologically more secure than other natural nourishments like meat, milk, eggs, poultry and sea food[10]. Variety of fresh fruits and vegetables are enriched with carbohydrates with pH value ranging from 7.0 to somewhat acidic and gives suitable growing environment to bacteria, yeasts and moulds [11, 12]. As compared to vegetables, fruits have better track record as wholesome food[8]. Citrus fruits are known for their phytochemicals which they generally use in their defence against parasites [13]. These fruits generally contain organic acids in quantities sufficient enough to give them a pH value of around 4.6 or lower than that. The acidic environment and the type of the acid found is the major reason for the growth of acid tolerant microflora in citrus fruits[14]. Fungus like *Saccharomyces cerevisiae* and Lactic acid bacterial species such as *Lactobacillus plantarum* is predominantly found in salted fermented fruit and vegetables[15, 16]. It is noteworthy that nutritional requirement of bacteria generally requires amino acids but variety of mutant *Lactobacilli* existing in nature have lost their requirements for an exogenous source of amino acids[17]. Lactic acid bacteria also produce exopolymeric substances that contribute to the texture of different food products[18]. These are structurally divided as homopolysaccharides and heteropolysaccharides based on their chemical composition.

The present study is focused on the micro flora found in *Citrus aurantifolia* (Common name:-Key Lime) which is a hybrid of *C. micrantha and C. medica* and belong to the plant family rutaceae. Some species of this family have originated in the area from North-East India to South-Western China[19]. The Indian state of Assam is one of the candidate origins of many *Citrus* species and home of different varieties of citrus specie. Unripe *Citrus aurantifolia* fruit has considerably low pH in it and rich in carbohydrate and trace elements which makes it as a potential source for the growth of LAB[20]. Traditionally local people of Assam preserve these fruit in clean jars and add salt to reduce spoilage. They are also used in medicinal purpose such as stomach ailments[21]. The quest for lactic acid bacteria in traditional food and their capability of producing exopolysaccharide will enhance sustainable bioprospecting and may open new doors of research in the field of food microbiology and may give answers on their interaction with living environment.

## Materials and Methods

### Sample collection

Sample of two months old fermented *Citrus aurantifolia* fruit was collected from the locality of Guwahati, Assam. The sample was collected using sterile procedure and brought to the laboratory immediately after collection for routine analysis. Small portion of the fruit was chopped manually with sterile cutter and the juice was extracted by mechanical squeezing till the considerable amount of extract was obtained.

### Isolation of Lactic acid bacteria

The extract so obtained was diluted into 10^−1^ dilution by adding 9 volumes of sterile distilled water. Stepwise dilution was done up to 10^−4^ dilution by taking 1 ml of well-vibrated suspension and it was then added into 9 ml water blank tubes. 100 μl from maximum dilutions were spread plated on the selective media. In the present study, MRS Agar media was used for isolation. 100 μl of the prepared sample extract was taken from the 10^−3^ and 10^−4^ dilution and was satirically pipette out in the agar plates. With the help of sterile L-shaped spreader it was thoroughly spread and after incubation, individual ropy or mucoid colonies were selected and transferred into sterile MRS broth medium. The cultures of the broth medium were then purified by streaking in MRS agar plates. The isolates were then examined for colony morphology, Catalase reaction and Gram stain[22, 23]. Only the Gram positive and catalase negative isolates were selcted from MRS broth medium. For preservation, the glycerol stocks of samples were prepared by mixing 0.5 ml of active cultures and 0.5 ml MRS medium including 40% sterile glycerol.

### Molecular identification

DNA extraction kit was used for bacteria DNA extraction (QIAGEN DNAeasy Kit (Qiagen; Hilden, Germany). An aliquot (1 μL) of genomic DNA extracted from each isolate was used as a template to amplify 16S rDNA gene. Polymerase chain reaction was performed in a Professional thermo cycler. The 16s rDNA gene was amplified using two bacterial specific primers: 16S F27, forward 5’…AGA GTT TGA TC(AC) TGG CTC AG…3’ and 16S R 1492, reverse 5’…TAC GG(CT) TAC CTT GTT ACG ACT T…3’. The PCR products were electrophoresed and visualized under UV light to determine their sizes and quality. The PCR products were then gel purified and sent to Eurofins Genomics India Pvt Ltd, Bangalore, for sequencing.

### Sequence search and submission

The 16S rDNA sequences obtained were compared with known 16S rDNA sequences at National Center for Biotechnology Information (NCBI) database using BLAST (Basic Local Alignment Search Tool) algorithm obtained from; http://www.ncbi.nlm.nih.gov/BLAST[24]. The sequence was submitted in GenBank (accession number) using sequin (http://www.ncbi.nlm.nih.gov/Sequin/index.html).

### Construction of phylogenetic tree

Phylogenetic relatedness of the isolated strain was investigated by collecting 16s rRNA gene sequences for different *Lactobacillus* species from NCBI database. The non-redundant dataset of 450 sequences was considered for the phylogenetic analysis. Multiple sequence alignment of the collected dataset was done by using MAFFT program [25] version 7.427. As the dataset contained more than four hundred sequences, PartTree based FFT-NS algorithm was used for the alignment. The resultant alignment file was saved for further phylogenomics analysis. The tree was constructed using Maximum Likelihood method based on the Tamura-Nei model with 500 bootstrap replicates. For this purpose, MEGA7[26] was used while iTOL web interface[27] was used for tree viewing and editing purpose.

### Production of exopolysaccharides

A simplified synthetic media was used for EPS production and characterization study adapted from Grobben GJ et al [28]. Batch fermentation studies were carried out at 37°C with pH of 6.4 for 24 hr. After cultivation, broth cultures were centrifuged at 8000 rpm for 10 min at 4°C. Once the cells get separated, the supernatant was taken in an falcon tube. To the tube containing the supernatant, about twice the volume of chilled ethanol was added and kept overnight at 4°C to get the EPS in precipitated form. After the overnight incubation, the tubes were centrifuged at 10,000 rpm for 10 minutes at 4° C. The supernatant was discarded and the pellet was kept for drying.

### Partial Purification of EPS

EPS was partially purified by dialysis and subjected to lyophilisation (freeze-drying) for further analysis. Dialysis tubing (Cellulose membrane, 14,000 M.W. Cut-off) was prepared by adding lengths of the tubing to boiling water containing EDTA and sodium hydrogen carbonate and stirring for 10 minutes. 1.5 gram of crude EPS was dissolved in 15 ml of double distilled water. The dissolved EPS was packed with previously activated dialysis membrane. EPS was dialyzed against distilled water for 24 h at 37°C with four changes of water. Partially purified EPS obtained from dialysis was frozen at −20° C in deep freezer.

### Characterization of EPS

FT-IR spectra of partially purified EPS were recorded from 400 to 4000 wave number per centimetre. Spectra were recorded using Nicolet-is10 FT-IR spectrophotometer in the department of Applied Science, Gauhati University. Lyophilized EPS sample was taken and mixed with Potassium bromide with the help of a mortar and pestle. It was then subjected to a pelletizer to form a pellet. A graph was obtained showing the peaks for different functional groups.

### Analysis of produced EPS

Estimation of total carbohydrate was done by Phenol-sulphuric acid method [29] while reducing sugar estimation was done by DNS method[30]. Glucose was used as the standard Absorbance was recorded at 490 nm against concentration of glucose.

## Results

### Identification of Bacteria

Bacterial isolate was assigned to a genus based on key morphological characteristics. Furthermore Gram positive, catalase-negative and non-motile cells were presumptively identified as lactobacilli. The molecular identification method by 16S rRNA sequencing also coincided with the results obtained by conventional method of characterization. Each isolate was genotyped by sequencing of a 390 bp section of the 16S rRNA gene (Fig: 1). The 16SrRNA sequence was aligned and compared with other 16SrRNA gene sequences in the GenBank by using the NCBI Basic Local alignment search tools BLASTn program. The BLAST results of the sequence (accession number: KT198973) showed maximum of 89% identity with *Lactobacillus delbrueckii* species. In our phylogenetic analysis, majority of the *Lactobacillus* specie have been rightly placed in their proper evolutionary positions. The isolated strain was grouped with *Lactobacillus delbrueckii*. in a separate clade.

**Figure 1:**
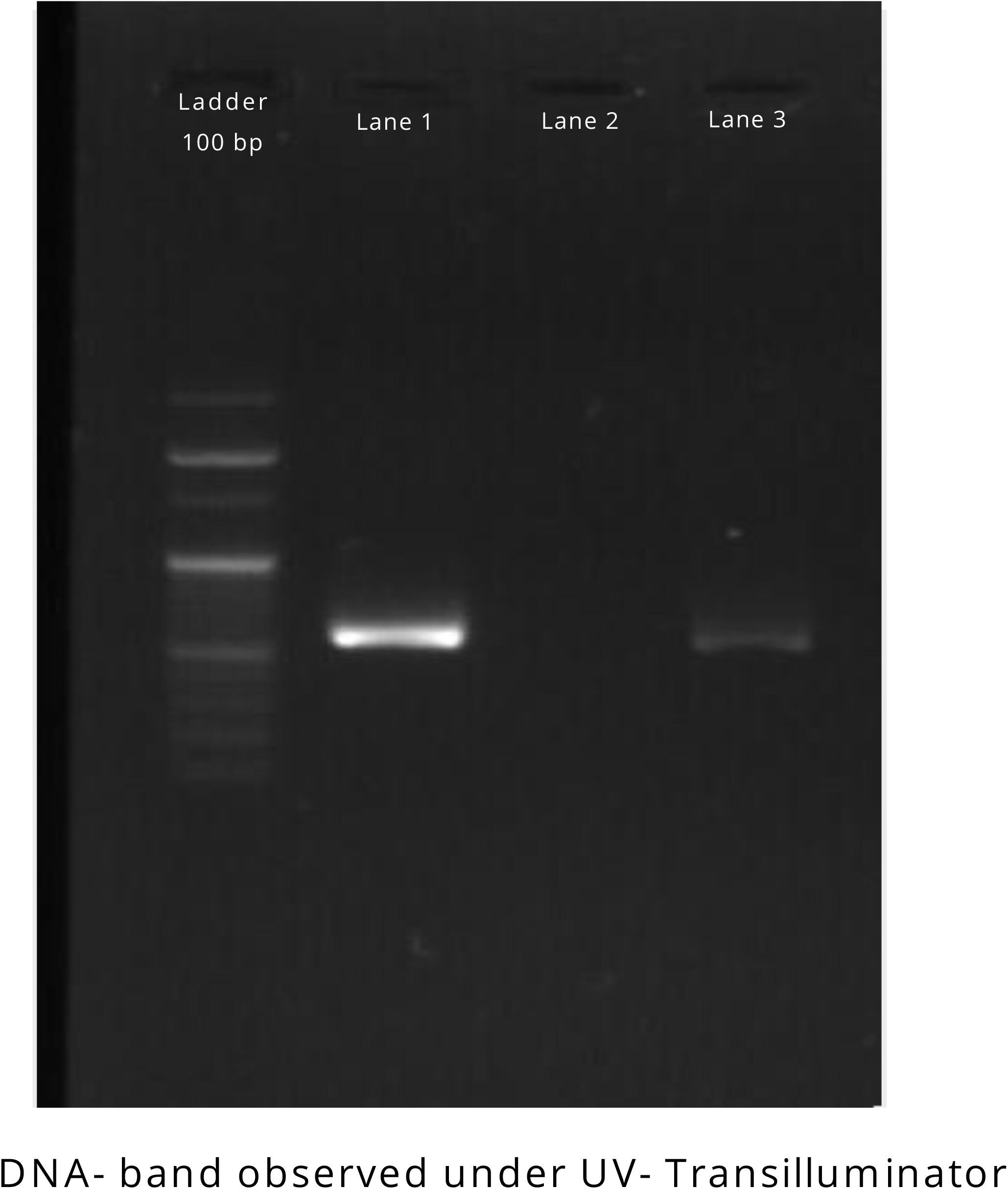
PCR amplification of 16 rRNA gene fragment. 100 bp DNA Ladder was used for approximate quantification. Lane 2 was the negative control (PCR water) while the other two lanes were the isolated bacterial sample.

### Production of EPS

After 24 hour of incubation in synthetic media, P^H^ of the media decreased from 6.4 to 5.6 due to the production of acid by the Lactic acid bacteria. Bacterial growth was monitored by absorbance measurement at 600 nm. The dry pellet obtained after centrifugation of fermented broth was the extracted EPS which was subjected to partial purification. Dry weight of the EPS obtained was determined.

### Estimation of total carbohydrate

Using the proposed method, the calibration curve was found to be linear in the range of 40-200mg/ml. A correlation coefficient of 0.9954 indicates good linearity between the concentration and absorbance. The total carbohydrate in EPS was calculated using regression equation obtained from the calibration curve (Fig: 2). The total polysaccharide content in EPS was found out to be 130.59 mg/ml.

**Figure 2:**
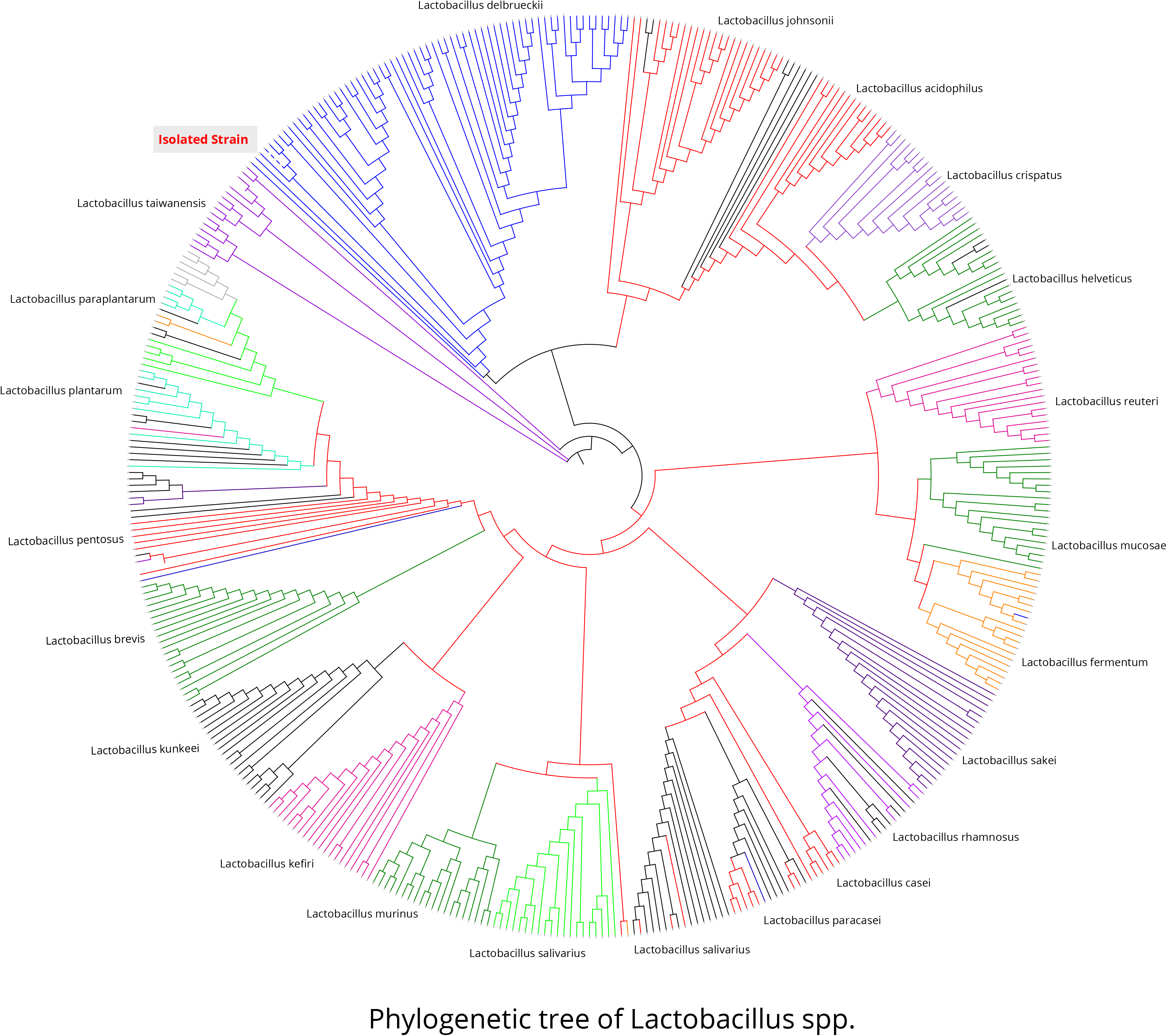
Maximum Likelihood based phylogenetic tree of *Lactobacillus* showing different species in their respective evolutionary position. The isolated bacterial strain was grouped with *Lactobacillus delbrueckii* in a separate clade.

### Estimation of reducing sugar by DNS method

The standard curve was found to be linear in the range of 40-160 mg/ml. The reducing sugar in EPS was calculated using regression equation obtained from the calibration curve and was found out to be 117.65 mg/ml (Fig: 3).

**Figure 3:**
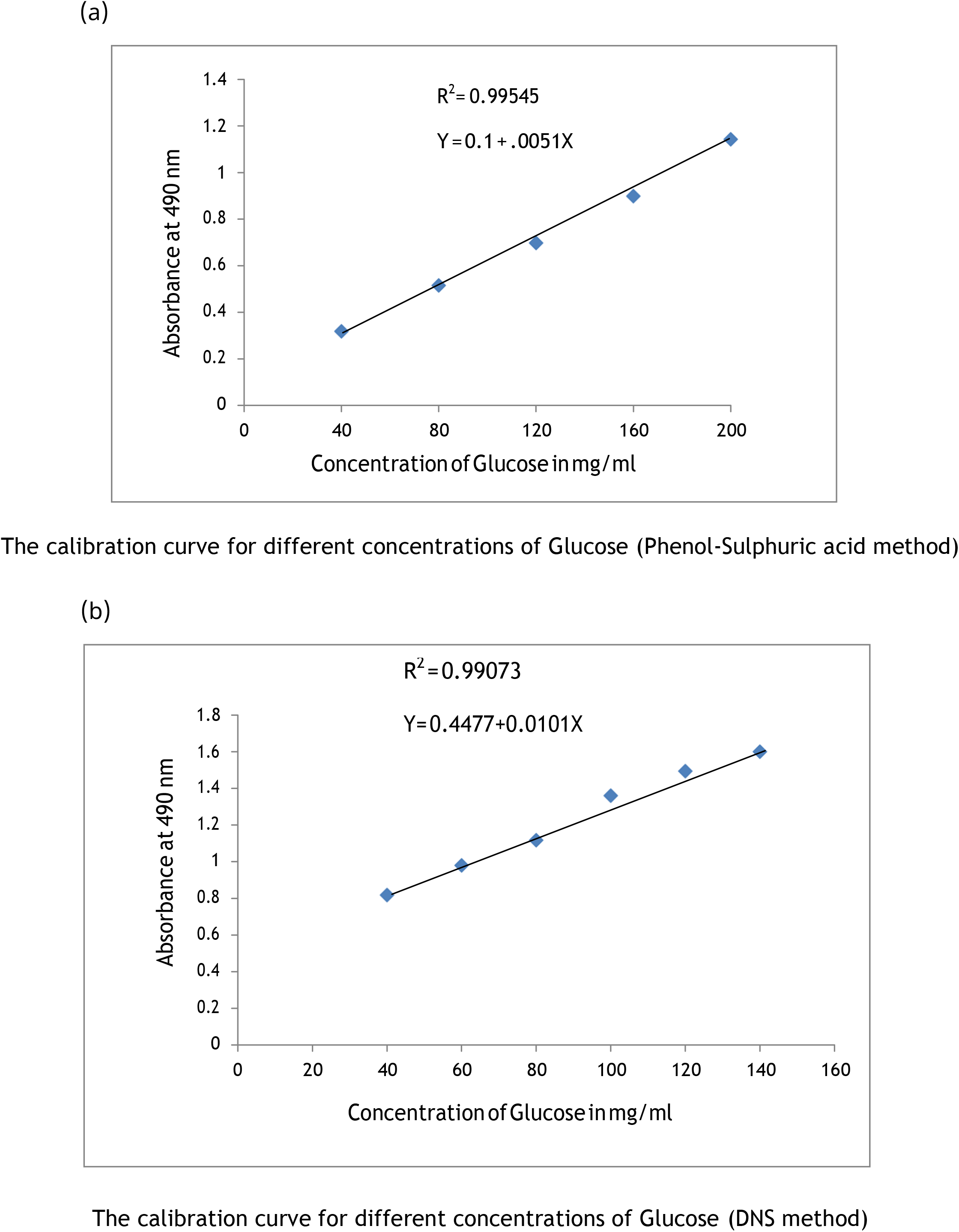
a) Determination of reducing sugars in partially purified exopolysaccharides by UV-visible spectrophotometry (DNS method). b) Estimation of total carbohydrate in partially purified exopolysaccharides by UV-visible spectrophotometry (Phenol sulphuric acid method)

### FTIR analysis

In the IR spectra a broad absorption band was observed at 3443c.m^−1^(Fig: 4). It indicates to the OH group vibrational stretching, usually shown by an open chain glucose molecule [31]. A medium absorption band was observed at 1644 c.m^−1^ which can be assigned as the stretching of the C=O bond[32]. Weak absorption peak were observed at 347 c.m^−1^. From this we can predict the presence of C-H bend of methyl group in the sample[33]. Absorption band at 1469 cm^−1^ is associated with aromatic and furan ring vibrations. A sharp absorption band at 1047 cm^−1^ corresponds to the presence of a sugar unit [34]. Some scattered small peaks were observed at 633,618 and 579 cm^−1^ which reveals the presence of alkane derivatives in the EPS[35].

**Figure 4:**
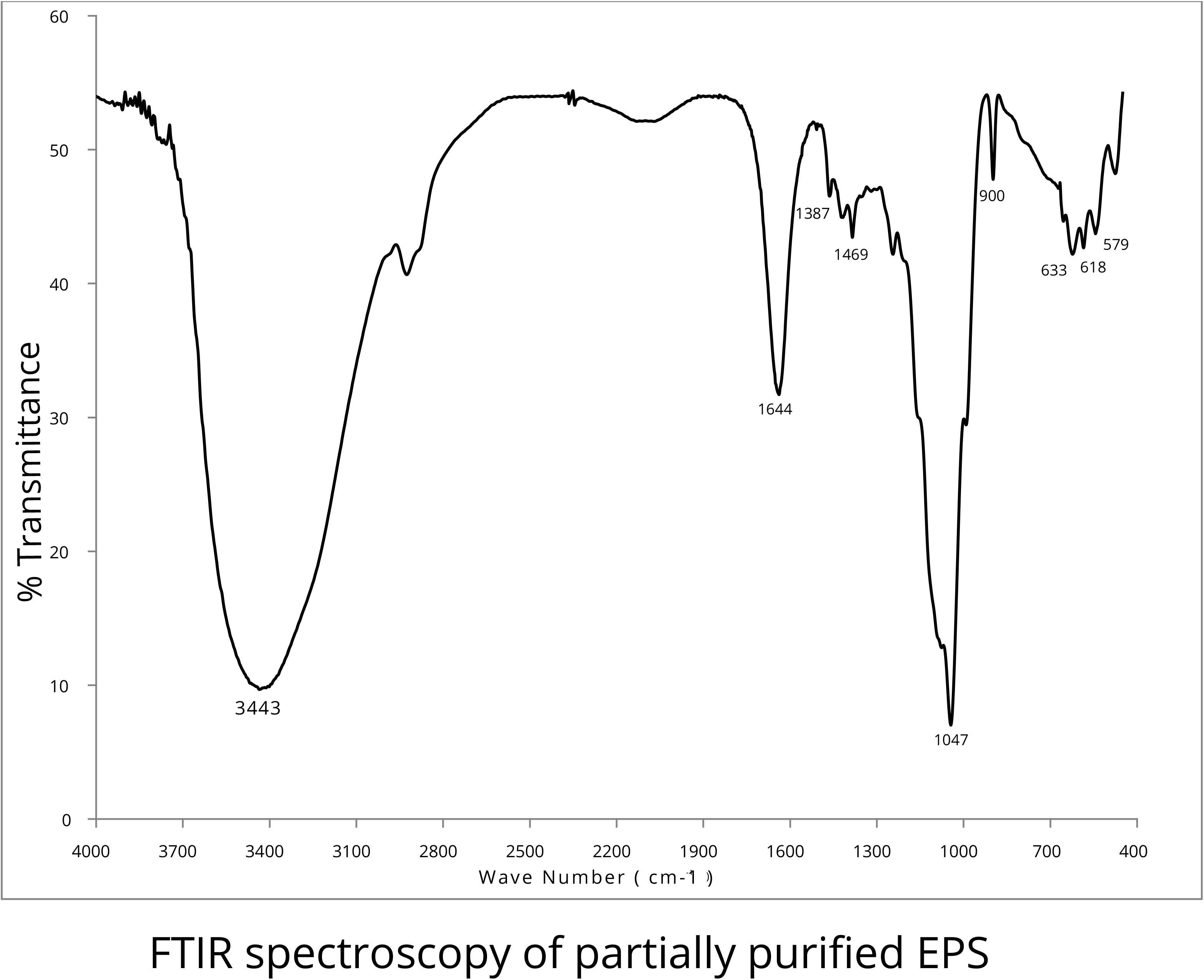
FTIR spectrum of partially purified EPS recorded using Nicolet-is10 FT-IR spectrophotometer.

### Discussion and future outlook

The outcome of 16s r-RNA gene sequencing of the isolated sample was quite interesting from the fact that sequence was similar with *Lactobacillus delbrueckii*, which is generally found in the starter cultures in the dairy industry; although the natural environment from which it has originated is not known for sure. To the best of our knowledge this is the first report of finding this sub species of LAB in citrus fruit product. Previous study reported the isolation of *L. delbrueckii* subsp. *bulgaricus* GLB44 along with its symbiont *Streptococcus thermophilus* from *Galanthus nivalis* (snowdrop flower) in Bulgaria[36]. Another strain of *Lactobacillus* delbrueckii subsp. delbrueckii TUA4408L was reported to be isolated from plant in Japan by Wachi et al[37]. Citrus extracts enhances the viability of *Lactobacillus delbrueckii*[38] due to enhance buffering capacity. However, it remains unclear how *L. bulgaricus* survive on plant products, and whether these bacteria are transferred from other materials. In order to differentiate the characteristics between commercial yogurt-starter strains and plant origin strain, further investigations need to be done. From the perspective of molecular characterization DNA-DNA hybridization, %G+C content, free fatty acid analysis and detailed biochemical and physiological characterization of the isolate will be done in the subsequent steps of this c feature of project. One of the characteristic features of *Lactobacillus delbrueckii* is the production of EPS and generally characterized by mucoid ropy colonies in the culture plate. Colony morphology of the studied strain prompt us to look for EPS producing capacity of the bacteria. The yield of EPS production by LAB is very less for homopolysaccharides and even lesser for the majority of heteropolysaccharides [39]. Furthermore, to remove the residual sugars of the culture media, dialysis of EPS is obligatory. EPS produced by the isolated bacterial strain in chemically defined medium was found to be 0.97 g/L that contained mostly sugar units. The obtained result could be used for designing optimal fermentation condition for over production of EPS.

### Conclusion

We have isolated a strain of Lactic acid bacteria from citrus fruit and identified it up to species level. EPS produced by the bacterial strain was purified by dialysis and chemical characterization of EPS was done by FTIR. The plant origin Lactic acid bacteria are of great interest to research and the finding will encourage the consumption of fermented *Citrus aurantifolia* juices as a probiotic food and EPS produced could have several applications in health and food industry.

## Acknowledgements

NB would like to thank to Department of Biotechnology, New Delhi; Department of Bioengineering and Technology, Gauhati University and Department of Horticulture, North East Hill University for providing support and assistance for the work.

## Conflict of Interest

The authors declare no conflict of interest and no competing financial interests.

## Author contributions

DB, AB designed the research; NB, AB performed text mining; NB carried out isolation and identification; AB performed computational and chemical analysis; NB, AB prepared the manuscript.

