## supplemental tables for "Isolation of a novel strain of Lactic Acid Bacteria from traditionally fermented common lime (*Citrus aurantifolia*) of Assam, India and analysis of exopolymeric substances produced by the strain"

**Supplementary:**

COMPOSITION OF EPS PRODUCTION MEDIUM

| Materials | g/L |
| --- | --- |
| Di-sodium hydrogen phosphate | 5.0 |
| Potassium dihydrogen phosphate | 6.0 |
| Tri-ammonium citrate | 2.0 |
| Sucrose | 50.0 |
| Magnesium sulphate | 1.0 |
| Trace elements solution | 10 ml |

TRACE ELEMENT SOLUTION COMPOSITION

| Materials | g/L |
| --- | --- |
| Ferrous sulphate heptahydrate | 5.0 |
| Manganese sulphate | 2.0 |
| Cobalt chloride | 1.0 |
| Zinc chloride | 1.0 |

TABLE OF ABSORBANCE VALUE FOR STANDARD AND SAMPLE IN PHENOL-SULPHURIC METHOD

| Sr. no. | Concentration (mg/ml) | Absorbance of Standard | Absorbance of the Sample | | | Mean |
|---|---|---|---|---|---|---|
| | | | Test 1 | Test 2 | Test 3 | |
| 1 | 40 | 0.319 | 0.768 | 0.764 | 0.766 | 0.766 |
| 2 | 80 | 0.516 | | | | |
| 3 | 120 | 0.698 | | | | |
| 4 | 160 | 0.899 | | | | |
| 5 | 200 | 1.39 | | | | |

TABLE OF ABSORBANCE VALUE FOR STANDARD AND SAMPLE IN DNS METHOD

| Sr. no. | Concentration (mg/ml) | Absorbance of Standard | Absorbance of the Sample | | | Mean |
|---|---|---|---|---|---|---|
| | | | Test 1 | Test 2 | Test 3 | |
| 1 | 40 | 0.819 | 1.639 | 1.640 | 1.631 | 1.636 |
| 2 | 60 | 1.132 | | | | |
| 3 | 80 | 1.268 | | | | |
| 4 | 100 | 1.526 | | | | |
| 5 | 120 | 1.701 | | | | |
| 6 | 140 | 1.891 | | | | |
